# Impact of Guanidine Hydrochloride on the shapes of Prothymosin-*α* and *α*-Synuclein is dramatically different

**DOI:** 10.1101/2024.10.01.616064

**Authors:** Zhenxing Liu, D. Thirumalai

## Abstract

The effects of Guanidine Hydrochloride (GdmCl) on two Intrinsically Disordered Proteins (IDPs) are investigated using simulations of the Self-Organized Polymer-IDP (SOP-IDP) model. The impact of GdmCl is taken into account using the Molecular Transfer Model(MTM). We show that, due to dramatic reduction in the stiffness of the highly charged Prothymosin-*α* (ProT*α*) with increasing concentration of GdmCl ([GdmCl]), the radius of gyration (*R*_*g*_) decreases sharply till about 1.0M. Above 1.0M, ProT*α* expands, caused by the swelling effect of GdmCl. In contrast, *R*_*g*_ of *α*-Synuclein (*α*Syn) swells as continuously as [GdmCl] increases, with most of the expansion occurring at concentrations less than 0.2M. Strikingly, the amplitude of the Small Angle X-ray Scattering (SAXS) profiles for ProT*α* increases till [GdmCl]≈ 1.0M and decreases beyond 1.0M. The [GdmCl]-dependent SAXS profiles for *α*Syn, which has a pronounced bump at small wave vector (*q* ∼ 0.5*nm*^−1^) at low [GdmCl] (≤ 0.2M), monotonically decrease at all values of [GdmCl]. The contrasting behavior predicted by the combination of MTM and SOP-IDP simulations may be qualitatively understood by modeling ProT*α* as a strongly charged polyelectrolyte with nearly uniform density of charges along the chain contour and *α*Syn as a nearly neutral polymer, except near the C-terminus where the uncompensated negatively charged residues are located. The precise predictions for the SAXS profiles as a function of [GdmCl] can be readily tested.

## Introduction

The discovery of Intrinsically Disordered Proteins (IDPs) is sometimes attributed to Pauling based on his theory of structure acquisition^1^ in antibodies in the presence of antigens. Fig. 1 in the above cited article clearly shows that the antibody undergoes a disorder-order transition only upon binding to the antigen, which is one of the features associated with IDPs (binding with a partner often leads to at least partial structuring). However, only through the pioneering studies of a few investigators in the mid-nineties the importance of IDPs in a variety of contexts was realized.^2–7^ It is now widely appreciated that IDPs are involved in a myriad of eukaryotic cellular functions. ^6,8^ Furthermore, certain IDPs form amyloid structures, which are associated often with neurodegenerative diseases. ^9^ The ability of IDPs to control a large number of functions is a consequence of their propensity to adopt a diverse ensemble of structures under physiological conditions. Unlike sequences that form ordered structures, IDP sequences have low complexity consisting of only a handful of amino acid residues. They are often enriched in charged and polar residues, which is illustrated clearly by the Uversky plot.^4^ Given the importance of IDPs, it is not surprising that intense efforts have been expended to understand their biochemical, biophysical and material properties using a variety of experimental and computational methods.

**Figure 1.**
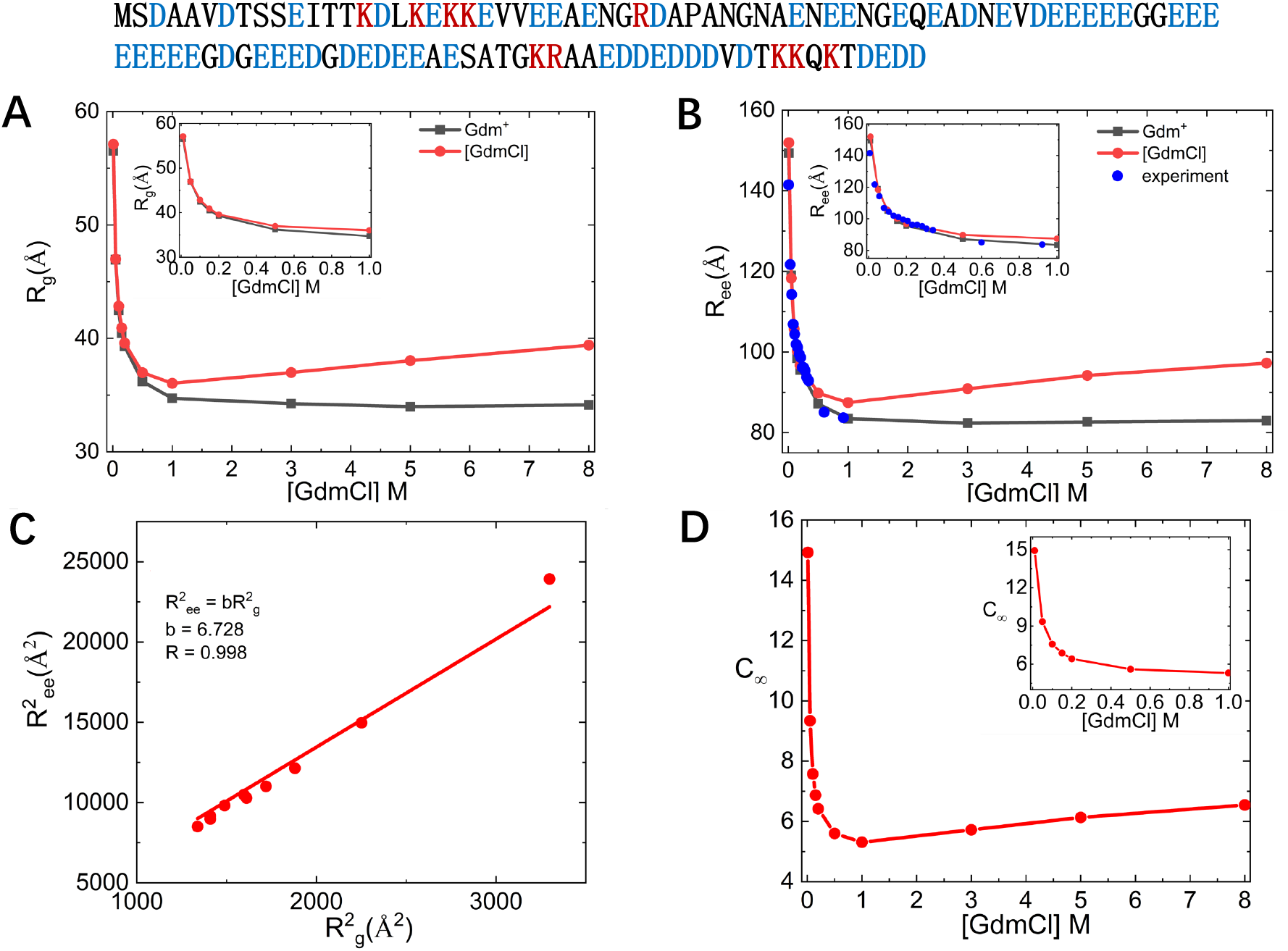
[GdmCl]-dependent sizes of Prothymosin-*α* (ProT*α*). The sequence of ProT*α* is shown at the top using one letter code for amino acids. The positively charged residues are in red and the negatively charged residues are in blue. (A) Radius of gyration *R*_*g*_ as a function of [GdmCl]. Black curve shows *R*_*g*_ calculated by accounting only for Gdm^+^ effects. Results in red are obtained using the MTM equation (Eq. 3). Inset is for *R*_*g*_ with [GdmCl] ranging from 0 from 1M. (B) Same as (A) except the changes in the end-to-end distance *R*_*ee*_ are plotted. Blue symbols are experimental results using KCl.^23^ In (A) and (B) the nonmonotonic dependence is obtained only if both the electrostatic and denaturation effects of GdmCl are taken into account through Eq. 3. (C) Linear correlation, with a correlation coefficient *R* = 0.998, between 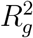 and 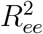 implies that the sizes at all values of [GdmCl] are determined only by one length scale. The fit is shown in the figure. (D) [GdmCl]-dependent *C*_*∞*_, which is a measure of the chain stiffness. Inset is for *C*_*∞*_ with [GdmCl] ranging from 0 from 1M.

The diversity of structures adopted by the IDPs enables them to respond to changes in environment, such as pH, temperature, and crowding. ^10–15^ For instance, conformations of IDPs with a large fraction of negatively charged residues should be responsive to changes in pH they would be protonated in acidic pH. In contrast, it is hard to anticipate the conformational changes in IDPs in the presence of denaturants, such as SCN^−^ (human metabolite) or Guanidine Hydrochloride (GdmCl), which is the focus of the present study. Although the denaturation mechanism of GdmCl in the unfolding globular proteins is reasonably well understood,^16^ it is more difficult to predict its action on IDPs. There are two main reasons. (1) GdmCl cannot induce cooperative transitions in the already unstructured IDPs in contrast to globular proteins, which undergo reversible transition from the folded to unfolded state. Therefore, one might be tempted to conclude that the shape changes induced by GdmCl would be minimal. (2) The precise sequence of the IDP could dictate the effects of GdmCl, which disassociates into weakly hydrated chaotropic ions Gdm^+^ and Cl^−^. Consider an IDP with *f*_+_ (*f*_−_) being the fraction of positive (negative) charge. The remaining residues could be polar or hydrophobic. Let *f*_−_ exceed *f*_+_. In this case, the favorable interaction of Gdm^+^ with the negatively charged residues^16^ could reduce the flexibility of the IDP. Such interactions are unlikely to be strong because of the charge density of Gdm^+^ ions is even less than Na^+^, for example. In the opposite limit (*f*_+_ *> f*_−_) the situation is less clear. (3) In addition to electrostatic interactions, GdmCl also has the effect of swelling any polypeptide chain, which has been amply demonstrated by its abilty to unfold globular proteins. The interplay between Gdm^+^ effects, in the charge dominated regime, and ability to swell the chain as the concentration of GdmCl increases, could elicit difficulty to predict responses in IDPs.

To elucidate the effects of GdmCl on the conformations of IDPs, we used the sequence-dependent transferable coarse-grained SOP-IDP model.^17,18^ The effects of GdmCl are taken into account using the Molecular Transfer Model (MTM).^19,20^ We performed Langevin dynamics simulations as a function of GdmCl concentration, [GdmCl]. The simulations are used to calculate equilibrium properties, such as the radius of gyration (*R*_*g*_), the end-to-end distance (*R*_*ee*_) and Small Angle X-ray Scattering (SAXS) profiles. The following are central results of our work. (1) The dependence of *R*_*g*_ and *R*_*ee*_ on [GdmCl] is dramatically different in the two IDPs. In Prothymosin-*α* (ProT*α*), both *R*_*g*_ and *R*_*ee*_ change non-monotonically,^21,22^ decreasing sharply at low [GdmCl], reaching a minimum at [GdmCl] *≈* 1.0M. Beyond 1.0M, ProT*α* swells. The precipitous decrease in *R*_*g*_ and *R*_*ee*_ is due to softening of the electrostatic repulsion between the negatively charged residues, which are roughly spread uniformly across ProT*α*. (2) In sharp contrast, *R*_*g*_ and *R*_*ee*_ in *α*-Synuclein (*α*Syn) increase rapidly at extremely low concentration of [GdmCl] (*≈* 0.2M) and then gradually expand beyond 0.2M.

The stiffness of the chain, calculated using the Flory characteristic ratio based on the *R*_*ee*_ as a function of [GdmCl], mirrors the dependence found in the variations of the sizes of the two IDPs. (4) We predict that the SAXS profiles change non-monotonically in ProT*α*. As [GdmCl] increases, the amplitudes of the SAXS profiles increases reaching a maximum at [GdmCl]*≈* 1.0M. Above 1,0M, there is a decrease. The SAXS profiles in *α*Syn have a pronounced bump at low [GdmCl] and the amplitudes decrease at higher [GdmCl], which is the generic behavior observed in IDPs. The contrasting predictions for the SAXS profiles in the presence of the GdmCl for the two IDPs are amenable to experimental test.

## Results

The dramatically different impact of GdmCl is illustrated using two IDPs with completely different sequence characteristics. To elucidate the effect of electrostatic interactions, we chose the 111-residue nuclear protein, ProT*α*, an IDP that regulates interactions with nucleosomes. The second IDP of choice is the 140-residue *α*-Synuclein (*α*Syn), which forms polymorphic amyloid fibrils and is involved in the Parkinson’s disease.

The fraction of positive (negative) charges is *f*_+_ = 0.09 (*f*_−_ = 0.49) in ProT*α*. The predominance of negative charges implies that electrostatic interactions between Gdm^+^ ions (with low charge density) should modulate the conformations of ProT*α*. The values of *f*_+_ = 0.11 and *f*_−_ = 0.17 in *α*Syn. Therefore, electrostatic interactions at neutral pH are likely to be less important but not negligible in *α*Syn. These arguments suggest that GdmCl should affect the conformations of the two IDPs differently, which we explore here using coarse-grained simulations.

### Prothymosin-*α*(ProT*α*)

#### *R*_*g*_ and *R*_*ee*_ change non-monotonically as a function of [GdmCl]

The size of the conformational ensemble is quantified using *R*_*g*_ and *R*_*ee*_, which are shown in Fig. 1 A and B. First, we ran simulations using Eq.2 by only considering the screening effect of Gdm^+^ ions (black symbols in Fig. 1A and B). The electrostatic screening gets more effective with increasing [GdmCl], which reduces the effective repulsion between the negatively charged residues that are nearly uniformly spread across ProT*α*. As a result, *R*_*g*_ of the polymer decreases sharply, reaching value of *≈* 35 Å at [GdmCl] around 1.0M (Fig. 1A). Gdm^+^ is expected to mimic monovalent ions, which could explain the precipitous drop in *R*_*g*_ up to [GdmCl] around 1.0M. If the denaturing effect of GdmCl is neglected *R*_*g*_ reaches a plateau at high concentration of Gdm^+^.

Similar behavior is found in the dependence of *R*_*ee*_ on [GdmCl] (Fig. 1B). The agreement between the simulations for *R*_*ee*_ (black symbols Fig. 1B) and experiments (blue dots in Fig. 1B)^23^ is excellent. When the concentration of Gdm^+^ increases beyond *∼* 1.0M, both *R*_*g*_ and *R*_*ee*_ reach a plateau (Fig. 1 A and B), because the screening effects are saturated.

We then performed simulations using Eq.3, which accounts for both the screening and swelling effects of GdmCl (red symbols in Fig. 1A and B). If the chain dimensions are determined solely by electrostatic effects of GdmCl, we would expect monotonic decrease in *R*_*g*_ and *R*_*ee*_ as [GdmCl] increases. In contrast, the simulations show that *R*_*g*_ and *R*_*ee*_ decrease non-monotonically with increasing [GdmCl] with a minimum at [GdmCl] *≈* 1.0M (see the red curves in Fig. 1 A and B). This result can be understood qualitatively by noting that the screening effect dominates up to 1.0M and thus the size of the conformation ensemble decreases. As [GdmCl] increases beyond 1.0M, the conformations are determined by the ability of GdmCl to expand polypeptide chains, leading to the increase of the size of ProT*α*. Although direct [GdmCl]-dependent measurements of *R*_*g*_ for ProT*α* are not readily available, we can estimate bounds from a couple of SAXS experiments done sometime ago. ^24,25^ The values of *R*_*g*_ varies between *≈* 38Å^24^ to *≈* 48 Å.^25^ Our predictions await future experiments.

Non-monotonic dependence of *R*_*g*_ as a function of [GdmCl] was reported for the N-terminal, C-terminal and full-length of ProT*α* in single molecule experiments^21,26^ by a careful analysis of the data using polymer theories. In addition, an interesting simulation study,^22^ which also used a combination of SOP-IDP and MTM, also found that *R*_*g*_ for ProT*α* decreases precipitously at small values of [GdmCl] and increases as [GdmCl] increases. Our results confirm these findings. Because of the preponderance of charged residues in ProT*α* (*f*_−_ *− f*_+_ = .40), it is tempting to rationalize the *R*_*g*_ and *R*_*ee*_ behavior in terms of polymer theories for polyampholytes or polyelectrolytes. ^21^ However, both sequence heterogeneity of ProT*α* and the chemical nature of GdmCl make it difficult to provide a clear molecular mechanism for the action of GdmCl (see the Discussion section).

#### *R*_*ee*_ and [GdmCl] dependent chain stiffness

The linear relation between 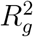 and 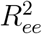 in Fig. 1C implies that globally there is only a single [GdmCl]-dependent length scale that determines the size.^18^ The data is well fit using 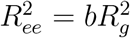 and *b ≈* 6.73. The value of *b* deviates from the predicted result, 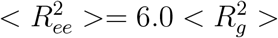, for an ideal freely jointed chain (FJC). Charge effects should change the stiffness of the polymer. In order to determine the stiffness, we exploit the dependence of *R*_*ee*_ on [GdmCl] to calculate,

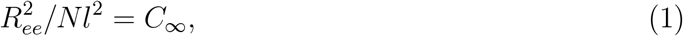

where *C*_*∞*_, the Flory characteristic ratio,^27^ is a measure of chain stiffness. The denominator in Eq. 1 is the size of an ideal chain with *l* = 3.8 Å, the distance between *C*_*α*_ atoms in the IDPs. The value of *C*_*∞*_ is unity for FJC. For worm-like chains (WLCs) near the rod limit, without volume exclusion, 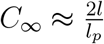 where *l*_*p*_ is the persistence length. The plot of the [GdmCl]-dependent *C*_*∞*_ (Fig. 1D) shows that the stiffness of the chain decreases first and then increases as [GdmCl] increases. At low [GdmCl], the value of *C*_*∞*_ is high (see the inset in (Fig. 1D)), reflecting the dominance of electrostatic interactions. At [GdmCl] *≈<* 1.0M, ProT*α* may be described as a stiff WLC. With increasing [GdmCl], *C*_*∞*_ decreases rapidly, as shown in Fig. 1D. The values at high [GdmCl] for ProT*α* are typically found in synthetic polymers.^27^

#### Small-Angle X-ray Scattering (SAXS) profile

We compare the experimental^24^ and simulated ([GdmCl] = 0.15M) SAXS profiles using the normalized Kratky plot in Fig. 2A. The data in grey are from experiment and the results in red are obtained by accounting for both the screening effect and denaturing effect of GdmCl. The data in blue corresponds to simulation results obtained by taking into account only the screening effect of Gdm^+^. Both the simulated results agree well with the experimental data, which again reinforces the efficacy of the SOP-IDP and the MTM in simulating the effect of GdmCl. At [GdmCl] = 0.15M, the conformational ensemble is largely controlled by charge interactions, which explains the coincidence of the results in blue and red in Fig. 2A.

**Figure 2.**
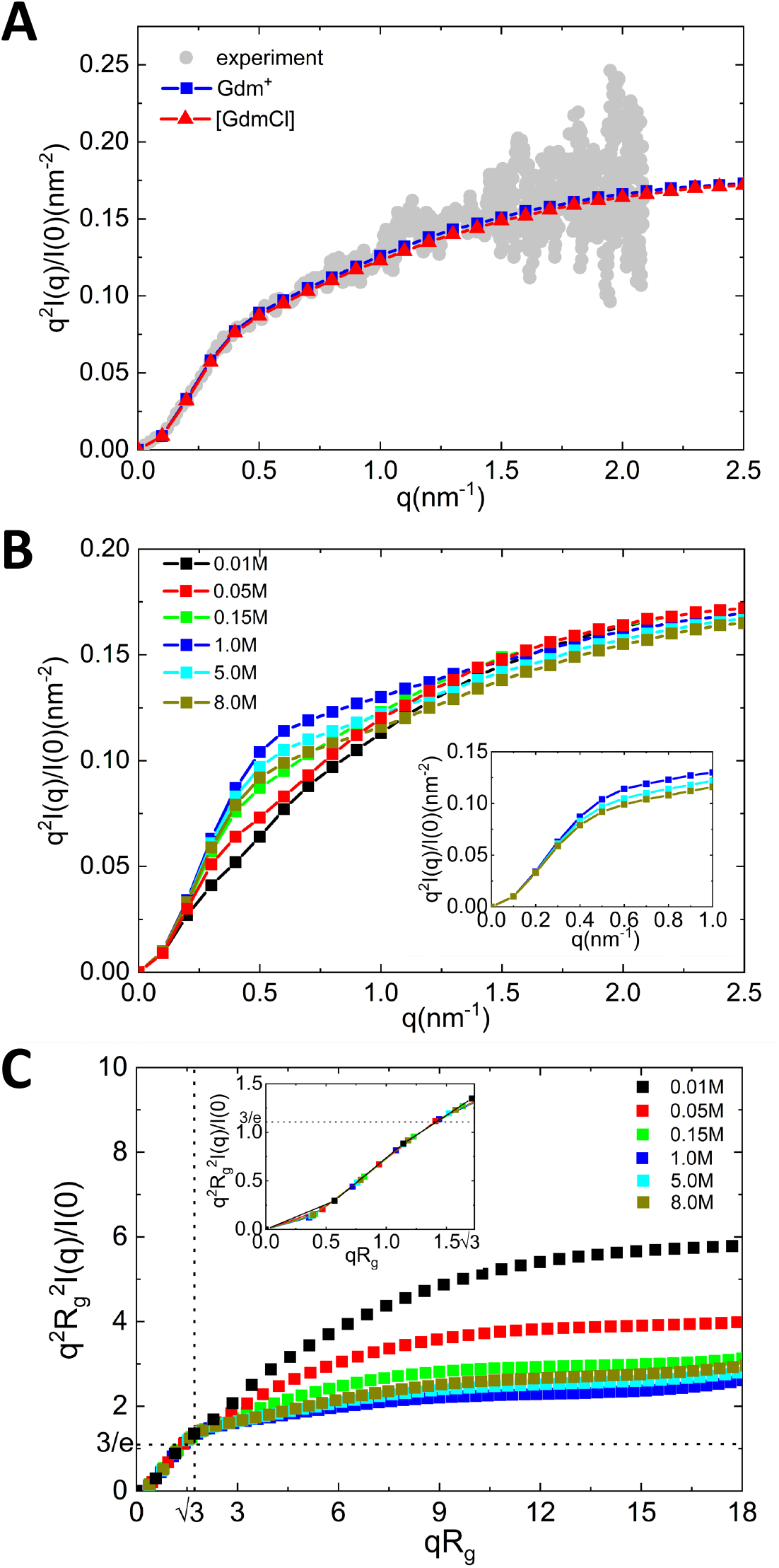
Kratky plots for ProT*α*. (A) Comparison between experimental and simulated ([GdmCl]=0.15M) SAXS intensity using normalized Kratky plots. (B) Normalized Kratky plots at different [GdmCl]. The inset shows the plots for [GdmCl]=1.0M, 5.0M, 8.0M when *q <*= 1.0*nm*^−1^. (C) Dimensionless Kratky plots at different [GdmCl]. Dotted lines are drawn at 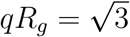 and (*qR*_*g*_)^2^*I*(*q*)*/I*(0) = 3*/e ≈* 1.104. The inset shows the plot for 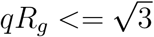.

In Fig. 2B, the dependence of the simulated SAXS intensity at different [GdmCl] using the normalized Kratky is plotted. At [GdmCl] greater than 1.0M there is a modest bump in *I*(*q*) for *q* less than 1*nm*^−1^. At higher [GdmCl], *I*(*q*) is featureless, which is characteristic of IDPs.^15,17,22^ However, it is striking that the SAXS profiles (Fig. 2B) increase at all values of the wave vector (*q*) till [GdmCl] *≈* 1.0M and decrease at higher concentrations (see the inset in Fig. 2C). The predicted non-monotonic changes at intermediate *q* implies that the distance distribution in ProT*α* varies in a non-trivial manner.

In Fig. 2C, we show the dimensionless Kratky plots to compare the simulated SAXS intensity at different [GdmCl]. The dotted lines are at 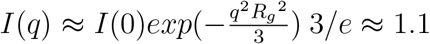and (*qR*_*g*_)^2^*I*(*q*)*/I*(0) = 3*/e ≈* 1.1. At low value of 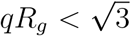, Guinier approximation gives 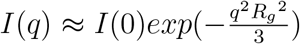 and thus the curves at different [GdmCl] overlap with each other. Although folded proteins have local maxima at the intersection point of the two dotted lines, in IDPs the scattering intensity increases after the intersection point.^28^ SAXS intensity of partially disordered proteins curves fall in between.^28^ Therefore, we can use the trends in the SAXS intensity beyond the intersection point as a guide to assess the extent of compactness of ProT*α*. Fig. 2C, shows that the conformational ensemble becomes more compact up to [GdmCl]=1.0M. Beyond [GdmCl]*>*1.0M, ProT*α* expands, which accords well with the [GdmCl]-dependent *R*_*g*_ and *R*_*ee*_ shown (Fig. 1).

#### Structure Factor

We extracted the Flory scaling exponent, *ν*, from the structure factor, *S*(*q*) (Eq.6) using, *S*(*q*) *∼ q*^−1*/ν*^ (Fig. 3A). Since the logarithmic plot for *S*(*q*) changes non-monotonically an small *q*, we choose an appropriate *q* range to calculate *ν*. For all values of [GdmCl], we used the *q* range indicated by the black line in Fig.3A. The extracted effective [GdmCl]-dependent *ν* is shown in Fig. 3B. When [GdmCl]=0.01M ProT*α* behaves as a stiff polyelectrolyte. The fit of *S*(*q*) *∼ q*^−1*/ν*^ in the *q* region indicated in Fig. 3A yields a value, *ν ≈* 0.85 (Fig. 3B). The high value of the effective *ν* shows that ProT*α* may be thought of worm-like chain that is close to the rod limit (*S*(*q*) *∼ q*^−1^, *ν* = 1.0). As [GdmCl] increases, ProT*α* is closer to a SAW limit with *S*(*q*) *∼ q*^−1*/ν*^, *ν ≈* 0.60). At [GdmCl]=1.0M, the effective Flory exponent has the smallest value, *ν ≈* 0.62. Similar dependence of *ν* on [GdmCl] was also observed for ProT*α* in experiments although the minimum *ν* occurs at a lower [GdmCl].^29^ The analysis of single molecule Förster resonance energy transfer (FRET) data^29^ was used to estimate *ν* = 0.67 at [GdmCl]=6.0M (orange square in Fig. 3C)), which is close to the simulation result. Based on a recent experiment (see Fig. 4F in^23^) which shows that *R*_*ee*_ at very low ionic concentration is on the order of *∼* 150 Å, we suspect that the value of *ν* could be close to our predictions, although this requires a more quantitative analysis.

**Figure 3.**
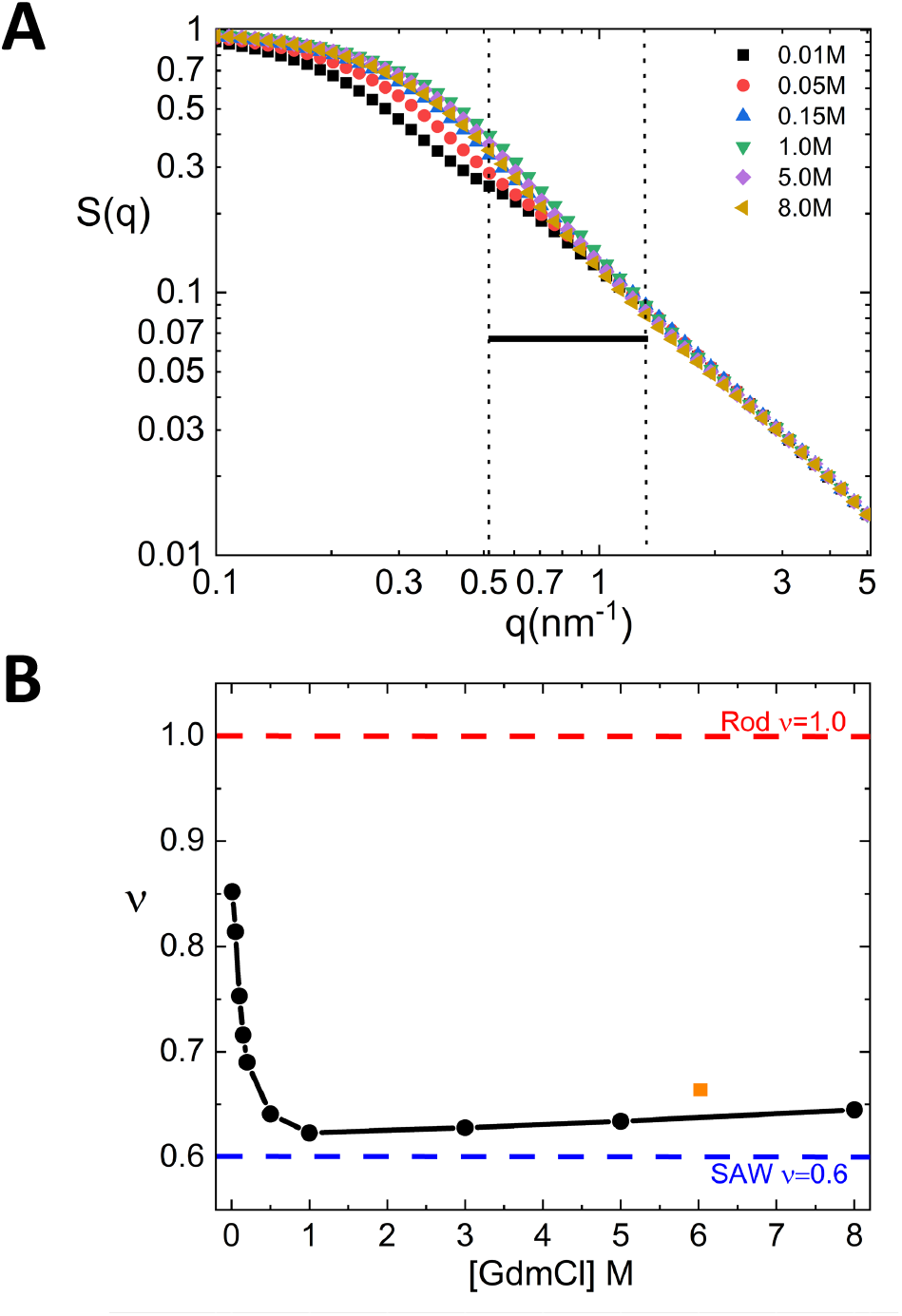
(A) Structure factor for ProT*α* when [GdmCl]=0.01, 0.05, 0.15, 1.0, 5.0, 8.0M. (B) [GdmCl]-dependent Flory scaling exponent *ν* for ProT*α*. For all [GdmCl], the data between two vertical dashed lines in (A) are used for fitting to get *ν*. The orange square is the experimental *ν* value.^21^

**Figure 4.**
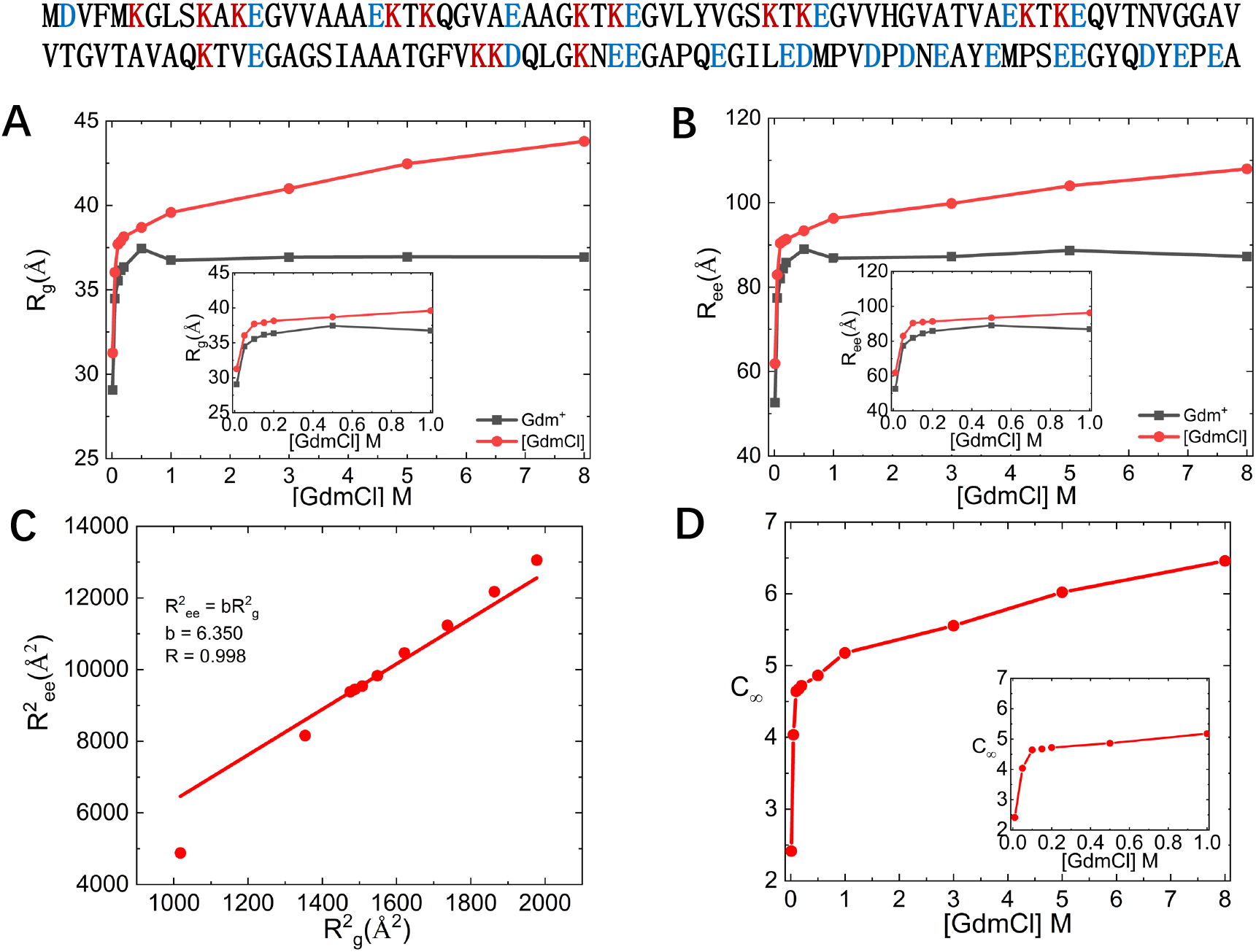
[GdmCl]-dependent sizes of *α*-Synuclein (*α*Syn). The sequence on top shows the positively charged (negatively) residues in red (blue).(A) Radius of gyration *R*_*g*_ as a function of [GdmCl]. Simulations results in black are obtained by taking into account only the ion effects of Gdm^+^. Results in red are obtained using the MTM model for GdmCl (Eq. 3). Inset shows *R*_*g*_ from 0 from 1M [GdmCl]. (B) Same as (A) except the changes in the end-to-end distance *R*_*ee*_ are plotted. (C) At all values of [GdmCl] 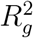 and 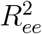 are linearly related. The fit is shown in the figure. (D) Dependence of the characteristic ratio, *C*_*∞*_, on [GdmCl]. Inset shows *C*_*∞*_ from 0 to 1M [GdmCl].

### *α*-Synuclein (*α*Syn)

#### [GdmCl]-dependent *R*_*g*_ and *R*_*ee*_

We then performed simulations to probe the effects of GdmCl on *α*Syn. Unlike ProT*α* in which negatively charged residues are (approximately) uniformly spread across the polymer (Fig. 1), in *α*Syn they are largely localized in the C-terminus. The N-terminus of *α*Syn is essentially neutral with the number of positive and negative charges being roughly equal. Therefore, we expect that *α*Syn should swell at very low [GdmCl] due to insufficient screening of the charges by GdmCl on *α*Syn. The [GdmCl]-dependent *R*_*g*_ and *R*_*ee*_ are shown in Fig. 4A and B (black symbols are obtained considering Gdm^+^ effect). In accord with the argument given above, *R*_*g*_ (Fig. 4A) and *R*_*ee*_ (Fig. 4B) increase sharply as [GdmCl] increases from 0.0 to *≈* 0.2M. Above this concentration, the swelling effect of GdmCl drives the increase in *R*_*g*_ and *R*_*ee*_ (Fig. 4A and B). Indeed, the increase in *R*_*g*_ and *R*_*ee*_ are clear in simulation results that account for both the screening and denaturing effects of GdmCl. The results, shown in red symbols in Fig. 4A and B, confirm that the denaturing effect results in the expansion of the polymer.

In Fig. 4C, the correlation between 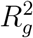 and 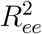 is shown. The calculated linear correlation is well fit using, 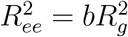 and *b ≈* 6.35. This again reinforces that in this IDP as well there is only one length scale (*R*_*g*_ or *R*_*ee*_) that determines the size. In Fig. 4D we plotted the [GdmCl]-dependent *C*_*∞*_, which shows that the stiffness of the chain increases monotonically as [GdmCl] increases. The *C*_*∞*_ values increase monotonically and non-linearly as [GdmCl] increases, which is qualitatively different from the behavior observed in ProT*α* (compare Fig. 1D and Fig. 4D).

#### Small-Angle X-ray Scattering (SAXS) profile

The experimental and simulated ([GmCl] = 0.15M) SAXS intensity using the normalized Kratky plot are compared in Fig. 5A. The data in gray are from experiment, the simulation result in red are obtained by taking into account both the screening effect and denaturing effect of GdmCl. The result in blue is calculated by considering the screening effect due to Gdm^+^ ion. The simulated results agree well with experiment. The denaturing effect (data in red in Fig. 5A) does not significantly change the result at [GdmCl]=0.15M because the screening effect is dominant, which is also reflected in the [GdmCl]-dependent *R*_*g*_ and *R*_*ee*_ in Figs. 4A and B.

**Figure 5.**
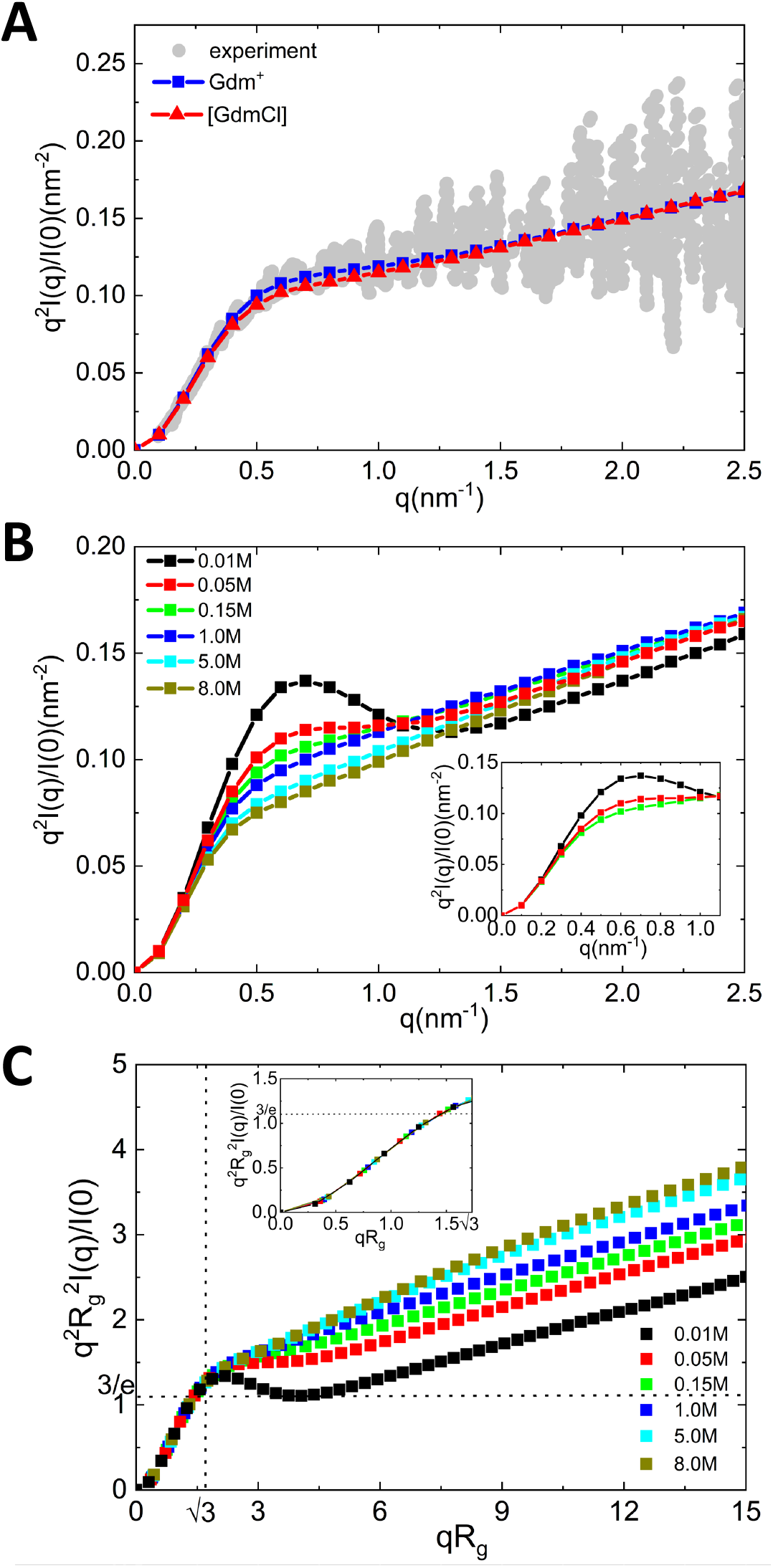
Kratky plots for *α*Syn. (A) Comparison between experimental and simulated ([GdmCl]=0.15M) SAXS intensity using normalized Kratky plots. (B) Normalized Kratky plots at different [GdmCl]. The inset shows the plots for [GdmCl]=0.01M, 0.05M, 0.15M when *q <*= 1.1*nm*^−1^. (C) Dimensionless Kratky plots at different [GdmCl]. Dotted lines represent 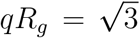 and (*qR*_*g*_)^2^ *I*(*q*)*/I*(0) = 3*/e ≈* 1.104. The inset plot is restricted to 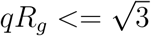.

In Fig. 5B, we present the predicted SAXS profiles at different values of [GdmCl] using the normalized Kratky representation. These predictions that there ought to a pronounced bump at *q ≈* 0.5 nm^−1^ at low [GdmCl] is experimentally testable. In Fig. 5C, we present the predicted SAXS profiles using the demensionless Kratky plot. At 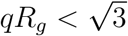, the Guinier approximation gives 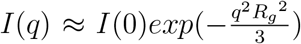, and the curves at different [GdmCl] coincide with each other (see the inset in Fig. 5C). In other words, 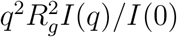 is a function of only *qR*_*g*_. From Fig. 5C, it is clear that as [GdmCl] increases, the conformational ensemble becomes more extended, which is confirmed by the [GdmCl]-dependent *R*_*g*_ and *R*_*ee*_ shown in Fig. 4A and B.

## Discussion

### Comparing the differences between ProT*α* and *α*Syn

The stark differences between the effects of GdmCl on ProT*α* and *α*Syn are illustrated in Fig. 1 and Fig. 4. ProT*α* undergoes substantial compaction as [GdmCl] increases from 0 to 1.0M, which could be due to softening of the electrostatic interactions caused by the interactions with Gdm^+^. In contrast, *α*Syn swells as [GdmCl] increases from 0M to about 0.2M. At higher [GdmCl] (*>* 1.0M for ProT*α* and *>* 0.2M for *α*Syn), the tendency of GdmCl to cause chain expansion is the dominant effect. These differing responses are due to substantial variations in the exact sequences between the two IDPs.

Despite differences at small values of [GdmCl], the linear relation between 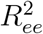 and 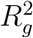 (Fig. 1C and Fig. 4C) shows that the sizes of both the IDPs are determined by a single length, as we showed previously^17,18^ for about 40 IDPs. This implies that polymer concepts could be used to establish certain universal relations. For instance, the ratio *ρ*_*eg*_ = *R*_*ee*_*/R*_*g*_ should have values that are characteristic of polymers. We find that *ρ*_*eg*_ for ProT*α* (*α*Syn) is 2.59 (2.52), which is within the range found for other IDPs.^18^

### On the polyampholyte (PA) and polyelectrolyte aspects of ProT*α*

Like ProT*α*, the monomers in polyampholytes (PAs) carry both positive and negative charges. This has inspired the use of analytical theory for random PA ^30^ to provide insights into the changes in the sizes of ProT*α* as [GdmCl] is varied. In an important study early,^21^ the dependence of *R*_*g*_ on [GdmCl] was extracted by analyzing the measured single-molecule fluorescence resonance energy transfer (FRET) data. They found that theory for random PA^30^ combined with a binding model for GdmCl interaction with ProT*α* quantitatively explained the behavior of *R*_*g*_ versus [GdmCl] (see Fig. 4 in^21^). This is surprising because the PA theory^30^ was derived by averaging over an ensemble of random sequences with a monomer in each sequence containing a positive or negative charge with a certain probability. In contrast, the sequence in ProT*α* is fixed (or quenched). Interestingly, the Fig. S3 in^21^ shows that a theory for polyelectrolytes in poor solvents^31^ also explains the *R*_*g*_ dependence on [GdmCl].

In contrast to the analytic theories, our simulations using the quenched sequence and an approximate molecular description of GdmCl interaction with ProT*α*, also nearly quantitatively reproduce the behavior of *R*_*g*_ as a function of [GdmCl] (see also^22^). Two comments are pertinent. (1) The near quantitative agreement between simulations and experiments shows that the combination of the MTM and SOP-IDP could be used as a basis to probe the effects of GdmCl on the conformations of IDP. (2) More generally, the remarkable success of polymer based theories to describe the sizes of IDPs discussed here and elsewhere^17,32^ deserves further scrutiny.

### Conformational heterogeneity

The generic polymer behavior noted here at high [GdmCl] hides the heterogeneity in the structural ensemble explored by ProT*α* and *α*Syn. For instance, hierarchical clustering of ProT*α* shows that 35% of the conformations are unusually compact.^17^ Similarly, about 44% of structures of *α*Syn are also highly compact. It has recently been shown theoretically that the presence of highly compact structures (heterogeneous structures) in the dilute phase could play a role in the propensity of IDPs to phase separate.^33^ Therefore, it is necessary to elucidate the sequence and environmental (such as pH, denaturants, and crowding) effects on the conformations of IDPs.

## Conclusion

Despite the success of the approach used here in reproducing experimental results, our work has limitations that have to be resolved before a molecular understanding of the action of GdmCl is reached. (1) We have treated Gdm^+^ as a point charge, which is a drastic approximation. In actuality, the single positive charge is delocalized over the three amine groups, making Gdm^+^ a low charge density cation. This aspect and the size of Gdm^+^ have to be explicitly considered in simulations. (2) Although the Molecular Transfer Model has proven to be accurate in predicting the effects of detaturants, it does not yield structural information about the interaction with IDPs at the residue level, which could be important in identifying the sub-ensemble of conformations that participate in phase separation or binding to other biomolecules.

## Methods

### SOP-IDP model

Several coarse-grained models^17,34^ have been proposed to simulate various aspects of IDPs. Here, we used the coarse-grained (CG) Self-Organized Polymer model for IDP (SOP-IDP), which has been successful in recapitulating a number of properties of IDPs.^17,18^ With the exception of a few models, ^35,36^ the majority use C_*α*_ (referred to as one bead per residue in the IDP literature) representation of the polypeptide chain, which were in vogue nearly thirty years ago to simulate the folding of globular proteins. In the SOP-IDP model, each residue is represented by two beads with one located at the C_*α*_ position and the other at the center of mass of the side chain. The SOP-IDP energy function is

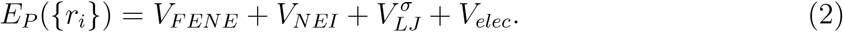

The detailed functional forms for *V*_*FENE*_, *V*_*NEI*_, 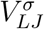, *V*_*elec*_ and the values of the parameters are given in the previous publications.^17,18^

The GdmCl has two main effects on the conformations of IDPs. As a weak acid, it disassociates into Gdm^+^, an ion with a low charge density. For simplicity, we treat Gdm^+^ as a monovalent ion that merely changes the Debye screening length as the concentration of GdmCl is altered. Thus, the screened Coulomb potential, *V*_*elec*_, used to treat the interaction between the charged residues on the IDP and Gdm^+^, becomes weaker as [GdmCl] increases. In addition, GdmCl acts as a potent denaturant, which we describe using the following Molecular Transfer Model (MTM).^19,20^

### Molecular Transfer Model (MTM)

At present, MTM is the only available computational method that accurately predicts the outcomes of experiments that use denaturants^19,37,38^ or pH as perturbation.^15,39^ In the MTM, whose theoretical basis is provided in a previous study,^20^ the effective free energy for a protein in the aqueous denaturant solution is given by,

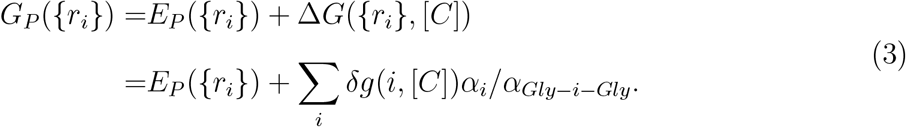

In Eq.3, Δ*G*(*{r*_*i*_*}*, [*C*]) is the free energy of transferring a given protein conformation from water to an aqueous denaturant solution at the concentration, [*C*]. The sum in the above equation is over all the beads, *δg*(*i*, [*C*]) is the transfer free energy of the *i*^*th*^ interaction center (C_*α*_ or the side chain of the amino acid), *α*_*i*_ is the solvent accessible surface area (SASA) of *i*, and *α*_*Gly−i−Gly*_ is the SASA of the interaction center *i* in the tripeptide *Gly − i − Gly*.

#### Langevin Dynamics Simulations

We assume that the dynamics of the protein is governed by the Langevin equation,

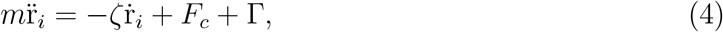

where *m* is the mass of a bead, *ζ* is the friction coefficient, *F*_*c*_ is the systematic force. We consider two cases. (i) We use *F*_*c*_ = *−∂E*_*P*_ (*{r*_*i*_*}*)*/∂r*_*i*_ (Eq. (2)) when treating the ionic effect, which alters the screening effect of GdmCl. (ii) The force is calculated as, *F*_*c*_ = *−∂G*_*P*_ ({*r*_*i*_})*/∂r*_*i*_ (Eq. (3)) if we take both the screening and denaturing effects of GdmCl into account. The random force Γ in Eq. 4 is assumed to have a white noise spectrum.

We performed Langevin simulations using a low friction coefficient, *ζ* = 0.05*m/τ*_*L*_ ^40^ and an integration time step *h* = 0.005*τ*_*L*_, where *τ*_*L*_ = 2*ps*.^41^ For each [GdmCl], we generated five trajectories, each with 10^8^ time steps. The equations of motions were integrated using the Verlet leap-frog algorithm.

#### Small-Angle X-ray Scattering (SAXS) Intensity

The SAXS intensity profile, *I*(*q*), is calculated using,

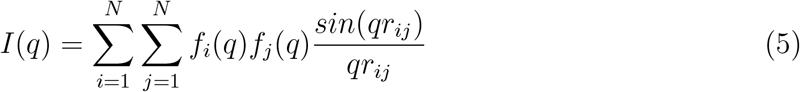

where *q* is the magnitude of the wave vector, *N* is the total number of beads in the IDP, and *r*_*ij*_ is the distance between the beads *i* and *j*. The values of *N* are *N* = 213 for ProT*α* and *N* = 262 for *α*Syn. The form factor *f*_*i*_(*q*) for bead *i* is taken from Table S3 in Ref.^42^

#### Structure Factor

The normalized structure factor, *S*(*q*), is given by,

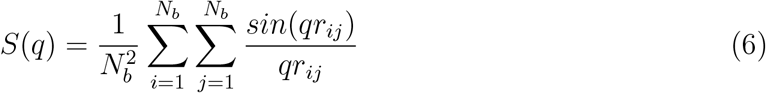

where *N*_*b*_ is the number of backbone beads, which is 111 for ProT*α* and 140 for *α*Syn.

## Acknowledgement

We are grateful to Julie Forman-Kay for a fruitful exchange on the history of IDPs. We thank Farkhad Maksudov for useful comments. ZL acknowledges financial support from the National Natural Science Foundation of China (11104015, 11735005). DT is grateful to the National Science Foundation (CHE 2320256), and the Collie-Welch Regents chair (F0019) for supporting this work.

## TOC Graphic

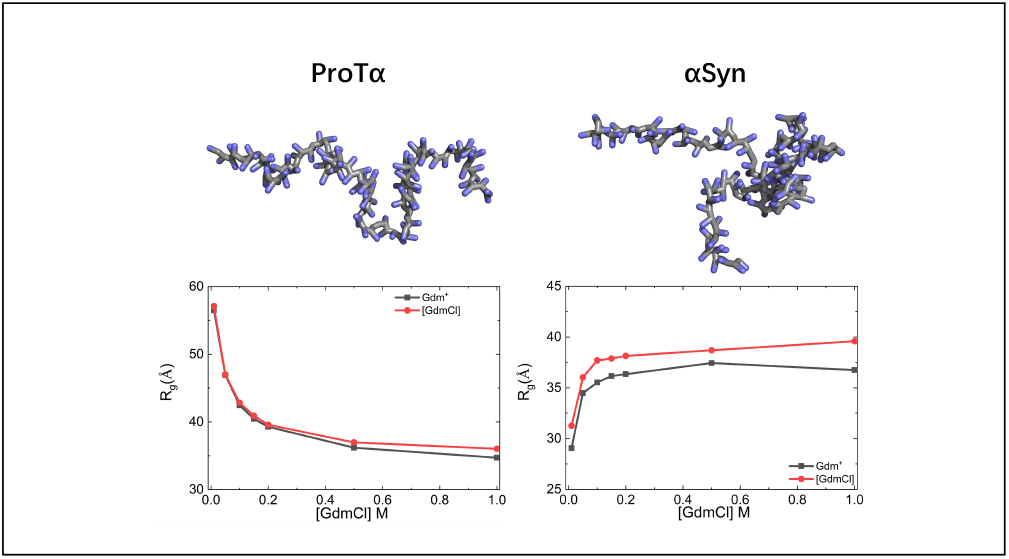

